# Long-term clonal analysis using stochastic models reveals heterogeneity and quiescence of hematopoietic stem cells

**DOI:** 10.1101/2025.09.24.677849

**Authors:** Yuri Garcia Vilela, Lars Thielecke, Artur C. Fassoni, Ingmar Glauche

## Abstract

Hematopoietic stem cells (HSCs) maintain lifelong production of blood by balancing self-renewal and differentiation. However, certain aspects of their divisional dynamics, namely the role of quiescence and the intrinsic heterogeneity of the HSC pool, are not completely understood. High-resolution clonal tracking provides a powerful resource to investigate such dynamics as the data captures patterns of clonal persistence, dilution and late clonal emergence. Here, we apply mechanistic mathematical modeling to longitudinal clonal data from non-human primates to explore structural requirements that underlie the observed dynamical patterns.

We show that models treating HSCs as a single, homogeneous population can explain the gradual loss of clonal diversity, but fail to reproduce clone size distributions and the long-term persistence of small and late-appearing clones. To address this, we propose a stochastic, two-compartment model in which HSCs transition reversibly between an actively cycling state and a quiescent, potentially niche-bound state. Compared to the simpler one-compartment model, this advanced framework provides a substantially improved fit for different metrics, consistently captures clone size distributions and explains the delayed activation and sustained coexistence of small and large clones.

These results provide quantitative evidence that heterogeneity within the HSC pool, particularly the existence of a reversible quiescent state, is critical to account for clonal aspects of long-term hematopoiesis. Our findings highlight how clonal data can uncover underlying regulatory mechanisms and supports a central role for niche-mediated HSC quiescence in maintaining stable and diverse blood production over time.

**Highlights:**

- Clonal data offers a robust approach to explore hematopoietic stem cell dynamics
- Homogeneous HSC population cannot reproduce observed long-term clonal patterns
- Niche-mediated quiescence improves model fit across different clonality metrics
- Model results suggest reversible transition between HSC quiescence and activation

## Introduction

Hematopoietic stem cells (HSCs) are crucial for sustained blood cell production throughout life by balancing self-renewal and differentiation. These rare, multipotent cells reside predominantly in the bone marrow and generate all blood lineages while at the same time maintaining their own stem cell pool. HSC activity is regulated by both intrinsic genetic and epigenetic mechanisms as well as external signals from the bone marrow microenvironment, also referred to as the stem cell niche [8, 9, 31, 29]. Few studies addressed the dynamical regulation of stem cell numbers and divisional activity with a particular focus on how the niche can regulate the balance between self-renewal and differentiation and thereby influence stem cell numbers [41, 13, 14, 27].

Clonal studies of HSC contributions rely on the identification and tracking of the direct offspring of individual stem cells, which can be achieved through artificial marking events, either in vitro [12, 30, 23, 6] or in vivo [28, 38], or by tracing naturally occurring mutations [21, 24]. By monitoring individual clones over time, such studies have elucidated key aspects of HSC self-renewal, long-term persistence and aging, shaping our understanding of hematopoiesis and its regulatory mechanisms [38, 23, 39, 24]. Clonal analysis also elucidated how HSCs and other progenitor cells contribute to blood production and how these differentiation pathways are challenged under stress conditions [3]. While gene therapy settings are a possible access to study post-transplantation clonality in humans, many aspects of dynamical engraftment and long-term polyclonality are still incompletely understood.

Non-human primates (NHPs) serve as an important model for studying clonal dynamics of HSCs in a more clinically relevant context [20, 32]. A remarkable contribution was the recent work by Radtke et al. [33] that followed individual HSC and progenitor clones over extended periods after autologous transplantation. The high-resolution tracking revealed that > 90% of long-term persisting HSCs contributed already to early hematopoietic recovery, challenging the assumption that only committed progenitors drive short-term hematopoiesis. Additionally, while their data showed a rapid loss of unique clones and the formation and expansion of long-term persisting clones, it also revealed the continued detection of new clones over time.

Radtke and colleagues use a mathematical, one-compartment model of HSC competition to quantitatively explain the manifestation of long-term persisting clones. This feature is known as clonal conversion and represents a hallmark of neutral competition in which stochastic processes lead to the extinction of some clones and the growth of others. The underlying model assumes that all HSCs are equally likely to divide or differentiate. Although the model explains the loss of clonal diversity between six months and four years post transplantation, it does not account for the delayed emergence of previously undetected clones even after four years. Moreover, the one-compartment model is insufficient to fully explain the distribution of clone sizes and the respective changes over time.

These observations led us to hypothesize that HSCs do not constitute a homogeneous pool of cells but rather a dynamic system in which cells can reversibly transition between an active state, characterized by division or differentiation, and a quiescent, potentially regenerative state. This interpretation follows a history of conceptual models that value this dynamic state transitions as an explicit feature of HSC organization, in which the hematopoietic niche plays a crucial role to support HSC quiescence [35, 41, 10, 25, 18]. One key piece of evidence supporting this view is that a homogeneous pool of continuously proliferating and differentiating cells fails to account for the outcomes of label dilution experiments, in particular the observation of long-term label retention [13]. The concept has also been applied to disease settings, such as leukemia, in which this tight regulation between HSCs and their regulating niches is disturbed, providing mutated clones with a competitive advantage [34, 11, 16, 15].

Here we propose a stochastic, two-compartment model of clonal hematopoiesis where HSCs can transition between an active cycling phase and a quiescent state within the niche. Testing this model against a simpler one-compartment model, we find that hematopoietic reconstitution involves not only stochastic fate decisions but also regulated transitions between quiescence and activation. Introducing a second compartment provides a mechanistic explanation for the persistence and delayed activation of certain clones, aligning better with long-term tracking data from the respective NHP models. The quantitative evidence for the existence of a quiescent compartment also suggests that interactions within the stem cell niche play a crucial role in long-term hematopoiesis.

## Methods

### Data

In a 2023 study, Radtke et al. [33] investigated clonal dynamics following autologous HSC transplantation after full-body irradiation in pigtail macaques, tracking the process at the clonal level from early recovery to long-term engraftment. Their high-density sampling approach uses genetic barcodes to produce a rich dataset containing temporal clonal information for two monkeys spanning over four years of follow-up.

Using this data available in the GitHub repository “ISA-Clone-tracking” [dataset][19], we extracted the clonal time courses of the two animals (Z13264 and Z14004). We restricted our attention to unsorted white blood cells (WBC) gathered from the peripheral blood, since this is the most complete and consistent data source over the whole experimental period (Supplementary Figure S1). We assume that the clonal composition of WBCs is a sufficient approximation of clonality at the HSC level.

### Clonality Metrics of the HSCs

From the extracted data, we derived three metrics reflecting different aspects of HSCs clonal dynamics (Figure 1):

- C*lonal diversity* is an approximation of the number of HSC clones present in the recipient animal at a given time point. Since there are no further transplantations inducing additional clones during the follow up, the number of clones progressively declines over time. Therefore, clonal diversity at a given time point refers to the number of remaining clones that are measured at this time point or any time point thereafter.
- S*ingle-time occurring (SO) clones* are those exclusively observed at a single time point. At a given time point, the *SO fraction* is defined as the number of SO clones divided by the total number of all observable clones. This metric approximates the abundance of small clones that are likely lost over time.
- The *relative clone size distribution* describes the pattern of clonal abundances observed in the marked population. At each time point, the read count of a clone is divided by the total number of reads to obtain its relative abundance. Clones are then grouped by abundance on a logarithmic scale, and the resulting histogram provides an empirical estimate of the overall clone size distribution.

**Figure 1.**
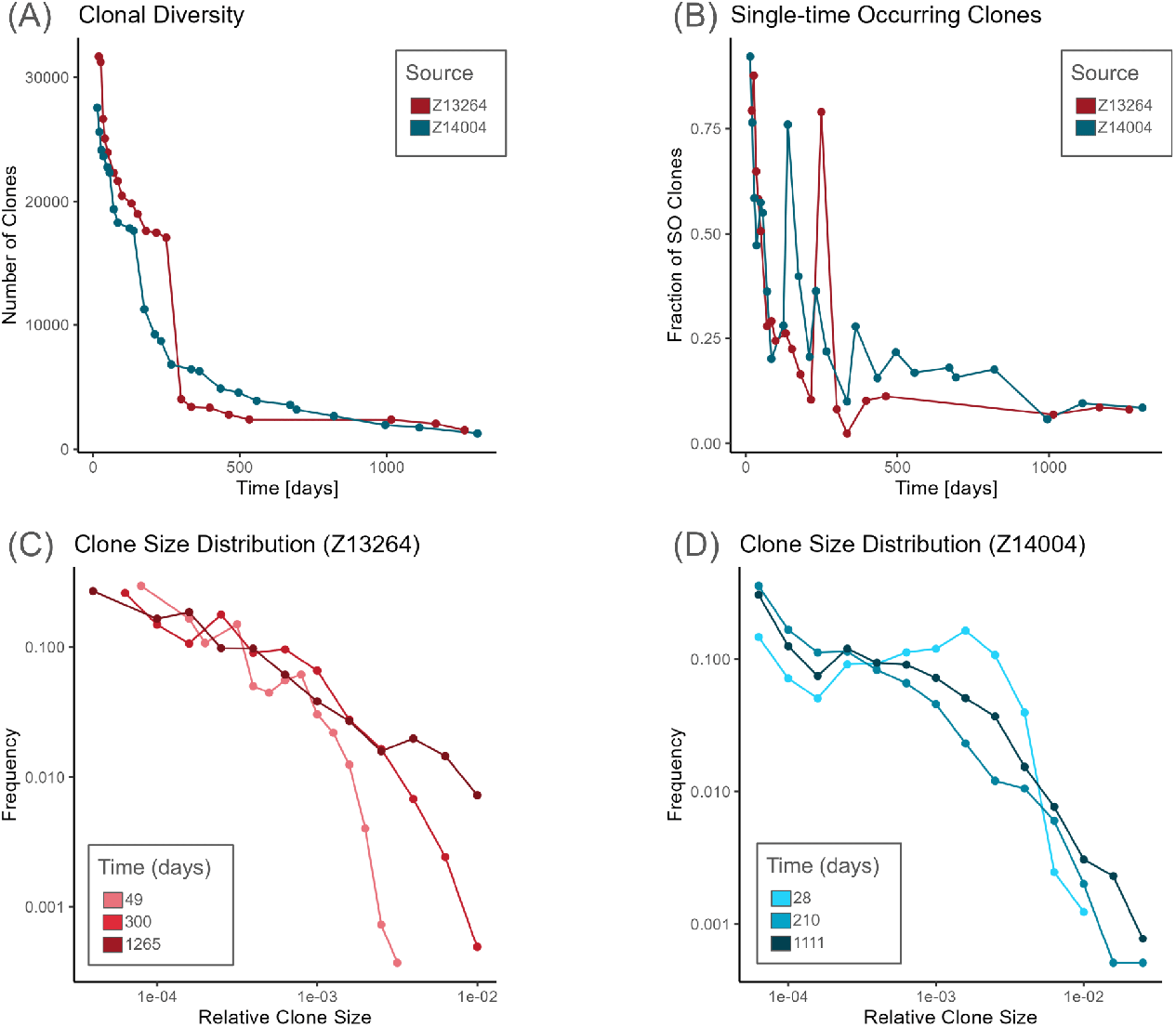
Metrics of HSC clonality: Metrics of HSC clonality, as derived from Radtke et al. [33] (A) Clone diversity, measured as the number of distinct clones detected over time. (B) Fraction of single-time occurring (SO) clones shown at each time point, i.e., the number of clones observed only at that time point divided by the total number of clones observed on that time point. (C-D) Clone size distribution for animals Z13264 and Z14004, respectively, shown as histograms of clone relative abundance (clone size divided by total of reads) on a log-log scale. Dots indicate observed data; lines connect observations for clarity. Colors denote animal id and sampling time as indicated in the legend.

Samples taken on day 249 for Z13264 and on day 138 for Z14004 are remarkable as they show an increased sequencing depth and detect a higher number of especially small clones. This manifests by the peak of SO clones in Figures 1B and the corresponding sharp drop in clone diversity in Figure 1A.

### Stochastic Models

A defining characteristic of hematopoietic stem cells (HSCs) is their dual ability to continuously generate differentiated blood cells while simultaneously sustaining their own population through self-renewal. Based on this principle, a simple stochastic model of HSC dynamics can be constructed, where cells either proliferate or differentiate at defined rates. As highlighted by Radtke and colleagues [33], a model based on these two mechanisms is sufficient to account for the progressive loss of clonal diversity over time. However, in order to account for other long-term phenomena such as label dilution kinetics [41, 2], it has been suggested that HSCs transition between different activity levels, namely an active state, in which they can proliferate and differentiate, and a predominantly quiescent state [35, 13, 25].

Here, we apply and quantitatively compare two different model structures, representing these distinct assumptions. In order to account for clonal identity, we formulate the model in a stochastic framework in which the cells can perform different actions with given propensities.

First, we consider a one-compartment model for the HSC pool, which is subdivided into n clonal subpopulations. Each clone proliferates at a maximal rate *p*_*A*_ and differentiates with rate *d*_*A*_. Proliferation is density-regulated by a logistic term that depends on the carrying capacity *k*_*A*_ and the total number of 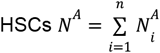, where 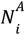 represents the abundance of clone i. At each time step, a cell can either proliferate, giving rise to two new HSCs of the same clone, or differentiate into a progenitor cell. The resulting proliferation and differentiation rates define the propensities of each event, and the likelihood of each event occurring is proportional to its corresponding propensity, as summarized in Table 1.

**Table 1.** Propensities for different cellular actions in the stochastic models. Each action occurs with a probability proportional to its propensity, which is determined by a base rate and the carrying capacity of the target compartment. The one-compartment model considers only active cells and actions of proliferation and differentiation, while the two-compartment model considers both cell states and all actions.

|  | Action | Propensity |
| --- | --- | --- |
| Active Cells | Proliferation | $p_A \left(1 - \frac{\sum A_i}{k_A}\right)$ |
| | Differentiation | $d_A$ |
| | Deactivation | $t_{AQ} \left(1 - \frac{\sum Q_i}{k_Q}\right)$ |
| Quiescent Cells | Activation | $t_{QA} \left(1 - \frac{\sum A_i}{k_A}\right)$ |

We also consider a two-compartment model that extends the one-compartment model by considering an additional compartment of quiescent HSCs, with clonal abundance 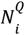 for each clone. The transition between the active and quiescent states is described by density-regulated transition rates *t*_*AQ*_ and *t*_*AQ*_ for deactivation and activation, respectively. Additionally, a carrying capacity *k*_*Q*_ for the quiescent compartment is considered. In this new model, the transition of active cells into the quiescent state is dependent on the base transition rate and the free-space in the target compartment, while quiescent cells have the ability to activate and return to the active compartment in an analogous way (Table 1). This model structure mirrors the one proposed by Hoffmann and colleagues [16], but is extended to accommodate an arbitrary number of clones with identical propensities.

A schematic representation of the second model is shown in Figure 2. It is straightforward to see that the first model is a special case of the second, obtained by removing activation and deactivation, i.e. setting 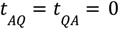. Furthermore, the one compartment model is equivalent to that proposed by Radtke and colleagues [33] when proliferation and differentiation have the same propensity.

**Figure 2.**
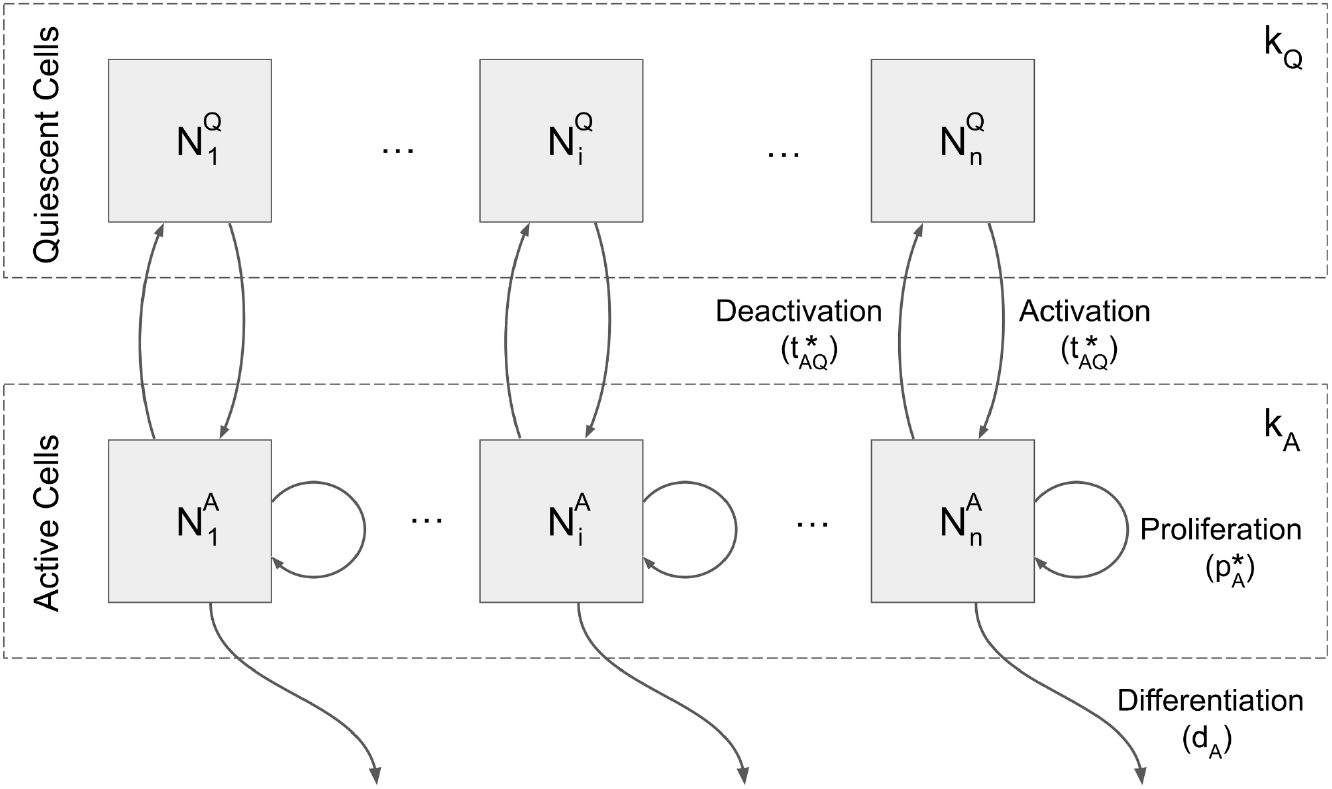
Two-compartment model of HSCs dynamics: Dashed boxes indicate the “active” and “quiescent” compartments with their respective carrying capacities, k_A_ and k_Q_, and one subpopulation for each clone represented as gray boxes. 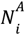 and 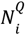 indicate the abundance of clone i in the active and quiescent compartments, respectively. Arrows indicate the flux of cells when the corresponding action is taken with propensities as defined in Table 1. A one-compartment model can be obtained by removing the quiescent compartment along with the actions that interact with it.

### Data for model fitting

The dataset from Radtke et al. [33] provides the clonal profile of two animals over a period of almost four years. We acknowledge that the early post-transplantation phase is characterized by dynamic fluctuations and transient contributions from short-lived, lineage-committed progenitors. In humans, it has been shown that these cells dominate hematopoiesis for several months, while long-term HSCs only gradually take over full hematopoietic functionality, typically after 6 to 12 months [3]. Likewise, studies in pigtailed macaques have shown that the hematopoietic system stabilizes over a period of 3 to 12 months before reaching homeostasis [32]. To avoid confounding short-term effects and capture more stable, long-term HSC-driven hematopoiesis, we focus our model analysis on time points beyond 360 days.

Moreover, as we are working with stochastic models, we aim to limit our analysis to more reliable measurements with lower intrinsic variability. For the SO clone fraction metric, we exclude the sample of animal Z13264 at day 532, as it only contains information on 17 cells. At such a small sample size even minor stochastic fluctuations across simulations tend to heavily impact this metric. Similarly, for the relative clone size distribution, we focus our fitting routine on small and medium-sized clones, with a relative size of around 10^-4^ and 10^-3^. In the data, these clones typically account for more than 85% of the total number of clones, providing a robust basis for model comparison. Evaluated on a logarithmic scale, these abundant clones are less influenced by stochastic fluctuations, whereas the rarer large clones (relative size 10^-3^∼10^-2^) display higher variability across simulations with the same parameters.

Furthermore, fitting of the clone size distribution is limited to two representative time points per animal (days 397 and 462 for Z13264, and 362 and 495 for Z14004). These have been selected based on high sequencing depth and temporal proximity to the beginning of the homeostatic phase. The limitation to two time points only is warranted to prevent an over-representation of this metric compared to the clonal diversity and the SO clones.

In Supplementary Table S1 we present a summary of the available data points and those used for model fitting.

### Simulation Routines and Optimization

In order to derive parameter configurations for each model class that optimally describe the experimental data, we minimize the squared sum of the distance between model and data on a logarithmic scale (residual sum of squares, RSS) across all metrics. To generate model predictions, we employ a customized R-routine that simulates clonal dynamics and the process of blood sampling at specified time points. Technically, the simulation proceeds as follows: given the number of clones and the rates described in the previous section, at each time-step we determine how many cells from each clone will perform a possible action, based on their propensities (Table 2). The carrying capacity is fixed at 10^5^ cells per compartment in the two-compartment setting, and at 2*10^5^ (active) cells in the single-compartment case (see Cosgrove et al. [7] and Mitchel et al. [24] for reference values in mice and men). The corresponding compartments are then updated accordingly. At time points corresponding to when blood was sampled in the *in-vivo* experiment, an equal number of cells is randomly chosen from the active compartment for clonal analysis. Tracking these sampled pools over time allowed us to derive temporal representations of clonal diversity, relative clone size distributions and the fraction of single-time occurring clones.

**Table 2.**
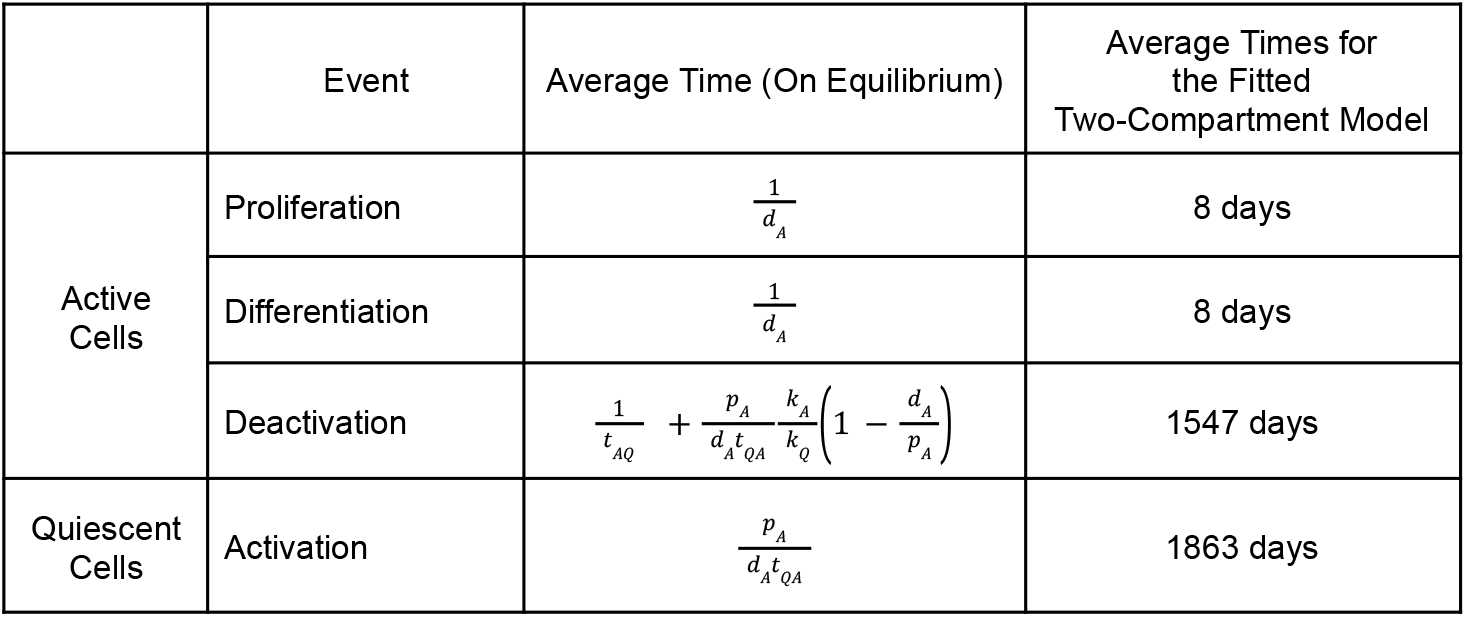
Analytical average time to cellular events at homeostatic equilibrium of the Hoffmann model [16]. Each value represents the expected time between two occurrences of a given action for a single cell, assuming the system has reached steady state. Expressions are derived as the inverse of the corresponding action rates in the deterministic ODE model described by Hoffmann et al. Average times for the best-fit parameters for the two-compartment model are shown in the third column.

For the optimization, we compare the model’s predictions with the actual data and quantify the difference in terms of the residuals for each data point (formally, the squared distance between model and data on logarithmic scale). Since the comparison is based on all three available metrics, we first normalize the residuals by the observed variation at each particular data point (obtained by repeatedly simulating with a previously defined parameter set), and then average the normalized residuals within each metric. Finally, the difference is calculated as the sum of the those residuals across all three metrics. This approach balances the contribution of each metric, ensuring that no single one disproportionally influences the fitting process.

For a given set of model rates, this approach returns a numerical value indicating the quality of the model’s fit to the data. We use this measure within a genetic algorithm [17] to explore the parameter space and identify optimal parameter sets for the initial number of clones and the event rates (p_A_, d_A_, t_AQ_, t_QA_ and p_Q_, see Supplementary Table S2). Technically, we use the GA package in R [37], which at each generation (iteration), generates a population of instances (i.e. set of parameters) which are evaluated with respect to the above quality measure. The best-performing instances (parameter sets within the population) are then used to generate the next generation, thereby gradually improving the fit over successive iterations. The process continues until convergence is reached, defined as the point where no further improvement in the fit is observed over several generations.

### Model Evaluation Metrics

For the optimization routine we use the normalized and averaged distance measure outlined above, which can be viewed as a generalized Residual Sum of Squares (RSS) on the log scale. For models with the same number of parameters, lower RSS values indicate better fit. To *compare different models* with different numbers of parameters, we additionally use the Akaike Information Criterion (AIC) [1] and the Bayesian Information Criterion (BIC) [36]. Both criteria incorporate penalties for model complexity based on the number of parameters, thereby helping to identify models that balance goodness-of-fit with parsimony. Lower AIC and BIC values indicate a better fit.

## Results

### Clonal convergence in a primate model

Tracking hematopoietic clonal contributions across species, including mice [12, 23, 6], non-human primates [32, 20], and humans [3], has revealed rich dynamic behaviors. Following clonal marking events, some clones expand while others contract, resulting in a gradual loss of clonal diversity over time. This phenomenon occurs even in the absence of selective advantages and reflects the inherent stochastic competition among individual cells. The term clonal conversion refers to this dynamic of reshaping the clonal landscape and the sequential loss of clones over time.

In a recently published dataset [33], Radtke and colleagues investigated the long-term behavior of genetically marked hematopoietic clones in pigtailed macaques (Figure 1). This comprehensive dataset allows deriving different diversity metrics, enabling detailed quantification of clone size distributions over extended time periods. It is interesting to see that clonal diversity drops rapidly in the early post-transplant phase, most likely as a result of the transient contributions from short-lived, lineage-committed progenitors (Figure 1A). However, on longer time scales, clonal diversity only decreases slowly. Similarly remarkable is the observation that even after several years post transplantation there is an occurrence of previously undetected clones that only occur once (Figure 1B). Changes in clone size distribution indicate that larger clones become more dominant while there is still a substantial fraction of smaller clones remaining (Figure 1C, D).

### Model based description of the data using a one-compartment model

To assess how well different mechanistic assumptions capture the outlined clonal dynamics, we systematically evaluated the performance of a one-versus a two-compartment model.

The one-compartment model represents a simplified view of hematopoiesis, where all HSCs remain in an homogeneous state, in which they continuously proliferate and differentiate. This model is equivalent to the one presented by Radtke et al. [33] and provides a reasonable fit for the overall measure of clonal diversity. However, when attempting to fit all metrics combined, the overall fit deteriorates significantly. Notably, in order to reproduce even a minimal fraction of SO clones beyond the one-year horizon (Figure 3B), the model requires an unrealistically large number of initial clones. This, in turn, leads to an overestimation of total clone numbers (Figure 3A) and a failure to reproduce the observed clone size distributions (Figure 3C–D). Even under these conditions, the predicted SO clone fraction remains well below experimental values, which consistently show the appearance of new clones even at later time points.

**Figure 3.**
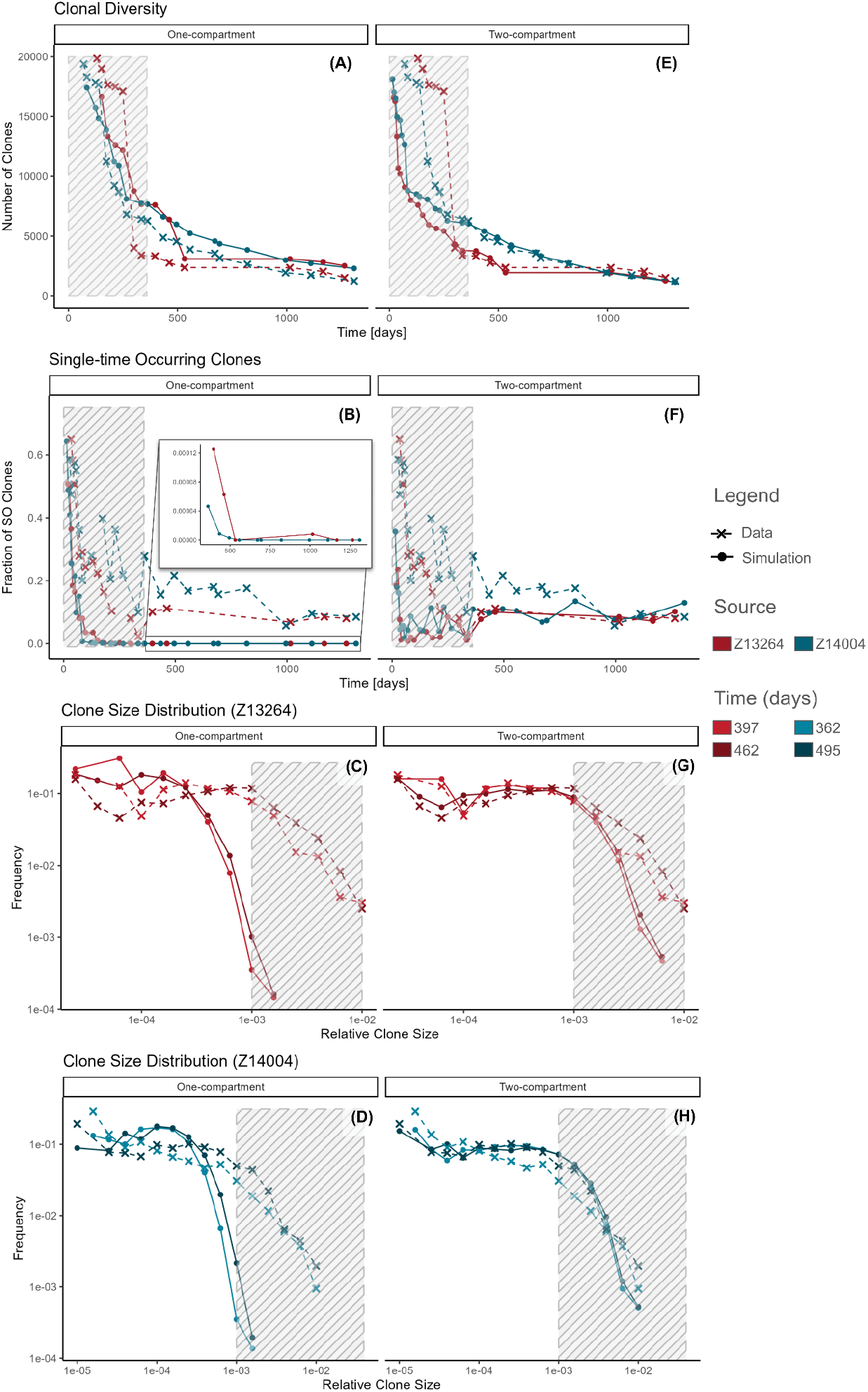
Model fits for the one vs two-compartment models: Best fit for the one (A-D) and two-compartment models (E-H) optimized with respect to the metrics derived from data by Radtke et al. [33]. Dots indicate model predictions with lines connecting observations for clarity; experimental data is indicated by **x** marks. Striped areas indicate data which is not used for fitting. Model parameters available in Supplementary Table 4.

Optimizing this model with a lower emphasis on the SO fraction yields visually improved fits, but still shows substantial discrepancies. Even the best fit achieved with this modification fails to capture the abundance of small clones, which appear on the left tail of the clone size distributions (Supplementary Figure S2). This limitation also affects the model-based approximation of the clonal diversity, leading to an underestimation of the overall number of clones.

Conceptually, these results highlight a key limitation of the one-compartment model: considering only one homogeneous HSC population, causes small clones to be more vulnerable to the competitive pressure from larger clones, making them unable to persist over long periods. This limitation suggests that additional dynamics are necessary to fully explain the observed clonal behavior.

### Model based description of the data using a two-compartment model

We propose that an intrinsic HSC heterogeneity, modeled as a transient quiescent state, improves the model predictions by explaining long-term clone persistence and the sporadic occurrence of small clones. To test this hypothesis, we introduce an additional compartment and the possibility to transition between active and quiescent states.

In the two-compartment model, HSCs are able to reversibly transition between an active state, where they proliferate and differentiate, and a quiescent state, where they remain proliferatively inactive. Analyzing the clonal profile generated by this model, we confirm that the inclusion of the quiescent state yields the expected results. Using this model, we were able to capture the continued emergence of SO clones throughout the entire sampling period (Figure 3F). Moreover, both the overall number of clones (Figure 3E) and their contributions to the HSC pool (Figure 3G and 3H) were in close agreement with the experimental data.

To quantify and compare the goodness of fit for each model, we evaluated the residual sum of squares (RSS), the Akaike information criterion (AIC) and the Bayesian information criterion (BIC). As shown in Supplementary Figure S3, the RSS for the two-compartment model is consistently lower compared to the one-compartment model across all evaluated metrics, confirming the visual intuition that the two-compartment model provides a significantly better fit to the data. Furthermore, both AIC and BIC scores are lower for the two-compartment model, indicating that even after penalizing the additional parameters, the two-compartment model remains better suited to describe the combined metrics (Figure 5 and Supplementary Figure S7).

We still notice an underestimation in the SO clone fraction during an intermediate period for animal Z14004, suggesting that additional mechanisms may be required to fully account for the increased presence of these small clones in this window. Despite this limitation, the two-compartment model successfully reproduces key features of the clonal data, reinforcing the idea that transitions between active and quiescent states are essential for shaping long-term persistence of clonal diversity, appearance and contribution patterns.

### Average Time to Events

The frequency of cellular events is a central aspect of biological dynamics, as it defines how often key processes such as division or differentiation occur. Fitting mechanistic models to empirical data enables the estimation of these frequencies or, equivalently, the inverse of it, which is the average time until an event occurs. In computational simulations, these quantities can be measured empirically by monitoring the time required for individual cells to undergo a specific event. As illustrated in Figure 4, the estimated average time for an event depends on the simulation time and stabilizes as it increases well beyond the critical time scale. This convergence behavior, typical of average-based estimations, highlights the importance of longer experiments to accurately capture the true temporal dynamics, especially for infrequent events.

**Figure 4.**
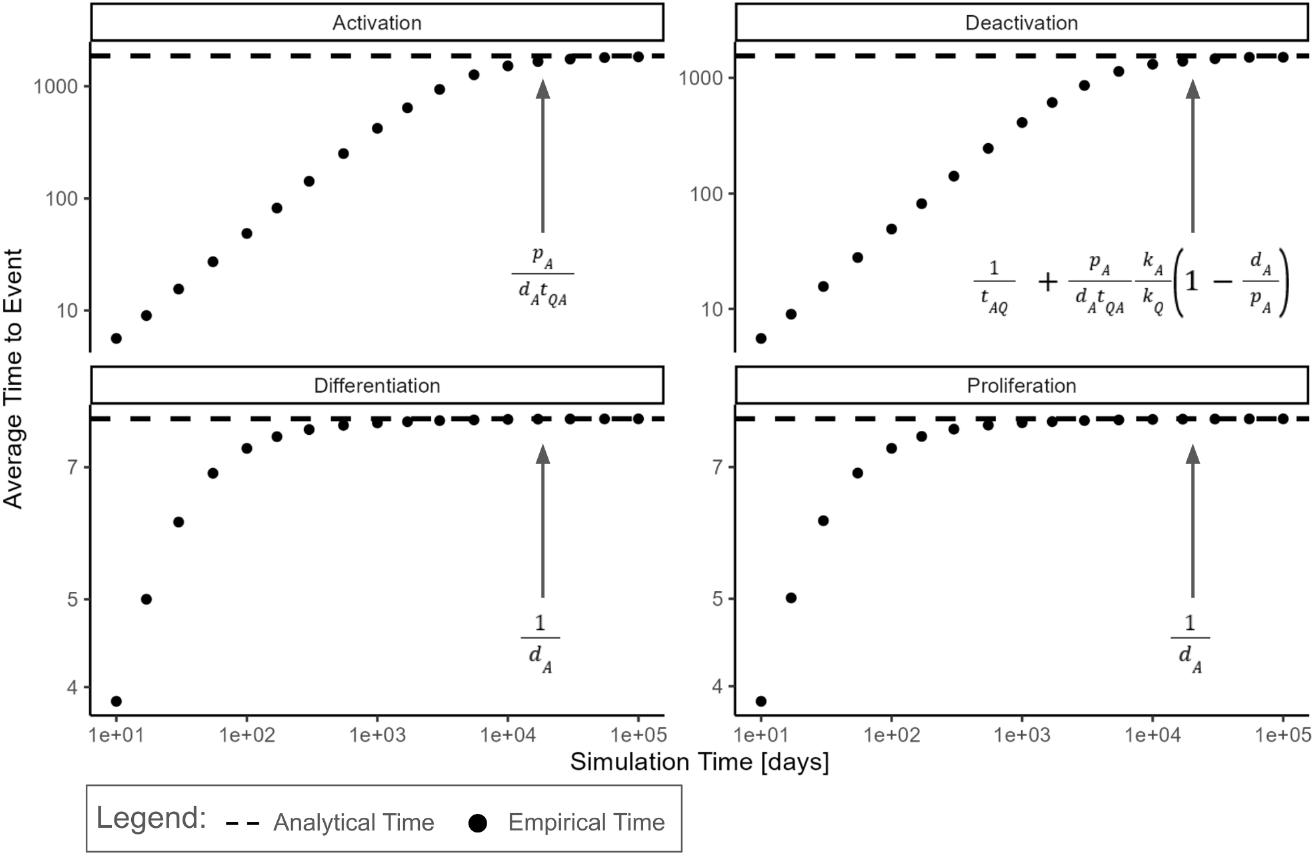
Average time to events: Empirical and analytical estimates for the average time to event at homeostatic equilibrium. Average times are estimated for the optimal parameter configuration obtained for the two-compartment model (Supplementary Table 4). Dots indicate the estimated average time between events recorded in the simulations at different time points. As the simulation period increases, the empirical estimates (dots) converge toward its analytically predicted counterpart (dashed lines).

Apart from this computational approach, one can also estimate these average times analytically. The stochastic implementation of the two-compartment model, can be seen as a generalization of the preceding ODE model [16] for the case of several clones with identical capabilities. In this setting, each cellular action (e.g., proliferation, differentiation, or state transition) occurs with a defined probability per unit time, proportionally to the rate used in the ODE formulation. At homeostasis, the expected time between successive events can be calculated as the inverse of its associated rate, with results summarized in Table 2. Figure 4 shows that the empirically derived average time between events converges toward its analytically predicted counterpart as the simulation time increases.

Evaluating the average time between events for the optimized two-compartment model, we observe that division-related processes (proliferation and differentiation) occur on a much faster timescale compared to state transitions (activation and deactivation). Namely, it takes on average one week for a cell to divide, while it takes on average several years for a cell to activate (Table 2). At first glance, this disparity might suggest that transitions can be neglected. However, omitting them effectively reduces the model to the one-compartment version, which cannot explain long-term persistence of small clones. It is interesting to note that transition times follow an exponential distribution (Supplementary Figure S4), implying that early transition events are still frequent despite long averages. While 50% of the cells transition within the first three and a half years, the fastest 25% of transitions occur within the first one and a half years (Supplementary Table S3). This characteristic of exponential distributions explains why such transitions, though rare in homeostatic situations, substantially influence the overall system behavior.

In summary, although state transitions occur much less frequently than division events, their presence is crucial for the long-term dynamics of the hematopoietic system.

### Different Mechanisms for HSC Proliferation

In the scope of the two-compartment model, the quiescent state is associated with niche regulation that maintains cells in a non-proliferative condition, while active cells divide independently of any previous actions (niche-independent proliferation). Recent work [26, 15], including our own, has suggested an alternative mechanism in which proliferation is directly linked to the transition from quiescence to activation (niche-dependent proliferation). To combine these views, a unified model has been proposed that incorporates both processes: in addition to random divisions of active cells, a fraction p_Q_ of cells exiting quiescence immediately undergo proliferation [15]. Notably, the two-compartment model discussed here can be perceived as a particular case of this unified model with p_Q_ = 0 (see Supplementary Figure S5). We raise the question whether this unified model is better suited to explain the long latency of hematopoietic clones after transplantation.

Incorporating the additional parameter p_Q_ in our fitting routine for the unified model, we observe that it achieves a fit similar to that of the earlier two-compartment model. Both reproduce key trends across all metrics with comparable accuracy (Supplementary Figure S6). This similarity is also reflected in overlapping ranges of the RSS (Figure 5A). Model selection criteria such as AIC and BIC (Figure 5B and Supplementary Figure S7) further reinforce this interpretation, showing only small differences between the two models. This suggests that niche-dependent proliferation may not be robustly identified from the shape of clonal dynamics under steady-state conditions, while a more prominent role in perturbed or stressed hematopoietic environments should not be excluded. However, comparing these metrics with the corresponding one-compartment model further supports the hypothesis that niche influence, whether through a quiescent compartment or via its role in proliferation, is pivotal to explain the observed dynamics.

**Figure 5.**
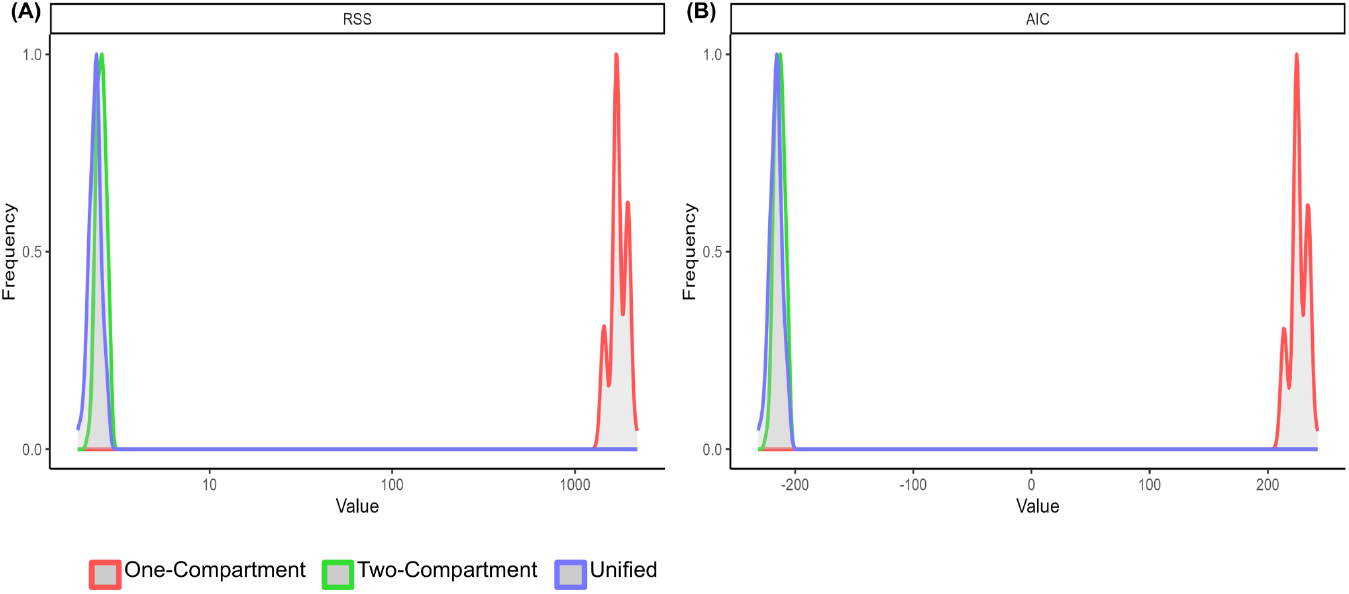
Model evaluation metrics: Distribution of Residual Sum of Squares (RSS) and Akaike Information Criterion (AIC) values from 100 simulations using the best-fit parameters (see Supplementary Table 4) of the one-compartment (red), two-compartment (green) and unified models (blue).

## Discussion

In this study, we employed high-resolution clonal tracking of hematopoietic cells in macaques to demonstrate that an additional layer of heterogeneity among HSCs is necessary to particularly explain the late emergence of smaller clones. Precisely, referring to a previously established concept, in which HSCs reversibly transit between a proliferatively inactive/quiescent state and an active, proliferative state, clearly outperforms simpler models that consider HSCs as a single, homogeneous population. Using a set of particular metrics to describe key features of the temporal clonal data set, we show that while a model incorporating only proliferation and differentiation can reproduce the gradual loss of clonal diversity, it fails to capture the observed distribution of clone sizes and entirely misses the long-term persistence of small clones. Introducing heterogeneity into the HSC population by distinguishing between actively cycling cells and niche-bound quiescent cells, yielded a remarkably improved fit to all metrics. Our results confirm that the addition of a quiescent compartment allows for a sustained coexistence of small and large clones and, in particular, the long-term emergence of single-time occurring clones across the observation period, which is in good agreement with experimental observations.

These findings highlight the importance of acknowledging a duality in the appearance of HSCs, to be either in an active or quiescent state. This concept has been long discussed on a theoretical level [22, 13], and is increasingly gaining recognition to explain biological data [41, 10, 40, 25, 18]. A prominent example arises from label dilution experiments [41, 25] where the biphasic decline of label-retaining HSCs, characterized by an initial rapid loss followed by a slower secondary phase, can only be explained by assuming that a substantial subset of HSCs exhibits markedly slower turnover dynamics.

It is an open conceptual question whether quiescence and activation in HSCs are truly two distinct and discrete states, or whether they are better understood as two extremes along a continuous spectrum of functional activity. In the latter view, individual HSCs may gradually shift their proliferative activity, potentially in response to external cues, but also to spatial and metabolic conditions. This would be in contrast to a sharp transition between binary states, as we consider it in our analysis. However, the two-state model offers conceptual clarity, is easier to represent in terms of mathematical or conceptual models, and also aligns well with experimentally accessible features such as flow cytometry based cell classification. In turn, it may only represent a simplified abstraction of a more complex biological reality.

While the division rates estimated for the two-compartment model are compatible with other data sources [41, 13, 5, 4, 3, 7], we notice that this model describes the observed data with fewer clones compared to the one-compartment model. It appears that the latter model structure tries to compensate for the rapid loss of clones by a higher initial clone number. Furthermore, while transitions between active and quiescent states emerge as a crucial mechanism to explain the data, their rate is much lower than the cell division rate. Still, the model fits clearly indicate that the slow but continuous flux of cells between compartments is a requirement for the steady renewal of the active pool, thereby accounting for the persistent emergence of new clones even at late sampling times.

Although our model provides a good overall fit to the data, some aspects remain unexplained. For instance, it cannot simultaneously reproduce the rapid loss of diversity observed shortly after transplantation and the slower dynamics that dominate during later phases. Moreover, while the model predicts the existence of single-occurring clones, it fails to reproduce their observed frequency at intermediate time points. These limitations suggest the need for more refined models that incorporate, for instance, heterogeneity in clonal capacities or context-dependent regulatory mechanisms. Importantly, such extensions can be naturally accommodated within the same two-compartment framework, offering a clear path forward for future work.

In summary, our analysis demonstrates that describing HSCs as a heterogeneous population comprising two complementary functional activity states provides a suitable explanation for the resulting clonal dynamics. It supports the notion of reversible HSC quiescence as a key element to understand long-term maintenance of clonal diversity. Our approach highlights the potential of using clonality data to explore the underlying mechanisms of HSC interaction and regulation, enabling the simultaneous evaluation of several different metrics in a unified framework and leading to more robust models.

## Author Contribution

**YGV:** Methodology, Software, Formal analysis, Investigation, Data Curation, Writing - Original Draft, Writing - Review & Editing, Visualization.

**LT:** Methodology, Data Curation, Writing - Review & Editing.

**ACF, IG:** Methodology, Investigation, Writing - Original Draft, Writing - Review & Editing.

## Acknowledgments

YGV was supported by the Coordenação de Aperfeiçoamento de Pessoal de Nível Superior - Brasil (CAPES) - Finance Code 001. LT was supported by the Deutsche. Forschungsgemeinschaft (DFG), grant GL 721/1-3 held by IG. ACF was supported by Alexander von Humboldt Foundation and Coordenação de Aperfeiçoamento de Pessoal de Nível Superior - Brasil (CAPES) - Finance Code 001, and partially supported by FAPEMIG RED-00133-21.

## Data and Code Availability

Data and code available on https://github.com/YGVilela/HSC.

