## Supplements for "Long-term clonal analysis using stochastic models reveals heterogeneity and quiescence of hematopoietic stem cells"

### Supplementary Figures and Tables

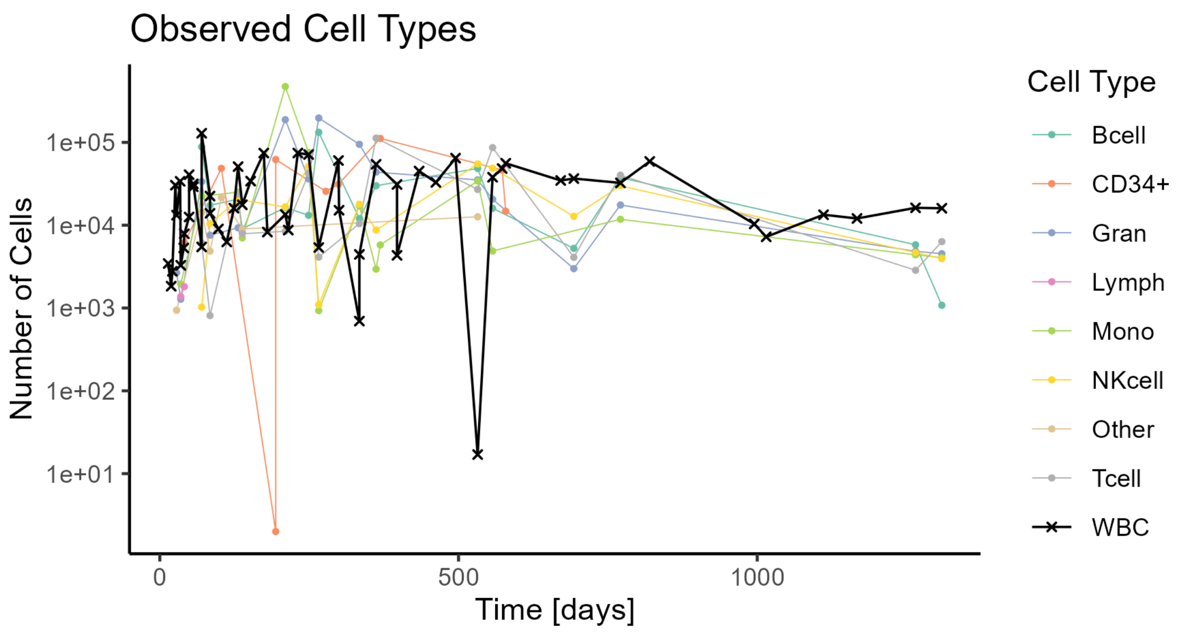


**Supplementary Figure S1:** Number of cells recovered for animals Z13264 and Z14004 from the data of Radtke et al. (2023) sorted by cell type. Points represent the observed number of cells other than white blood cells (WBC), represented by the **x** marks; lines connect observations for clarity. Colors denote the different cell types as indicated in the legend.

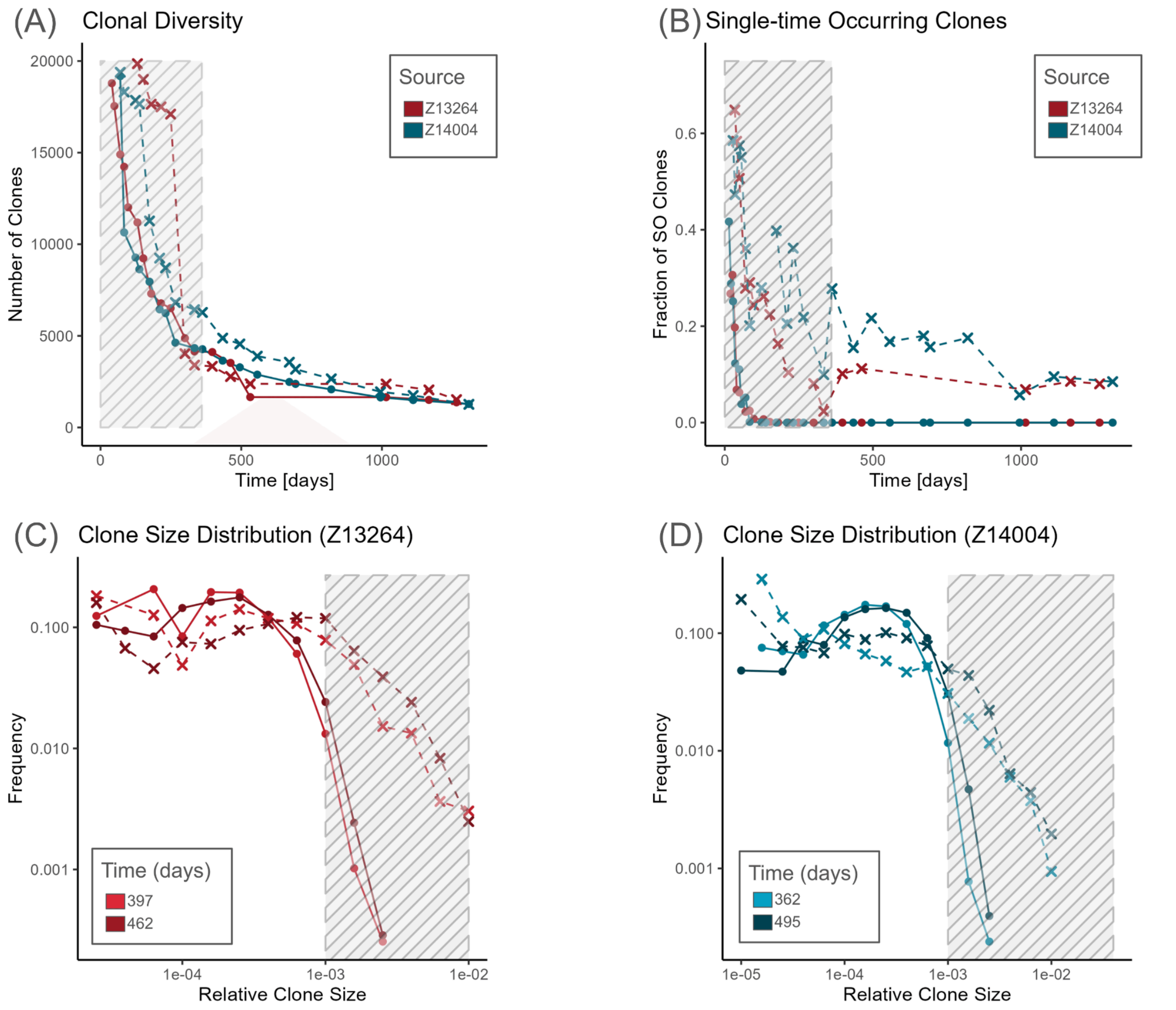


**Supplementary Figure S2:** Best fit of one-compartment model with SO clones residuals downweighted by a factor of 0.05 optimized with respect to the metrics derived from data by Radtke et al (2023). Dots indicate model predictions with lines connecting observations for clarity; experimental data is indicated by **x** marks. Striped areas indicate data which is not used for fitting.

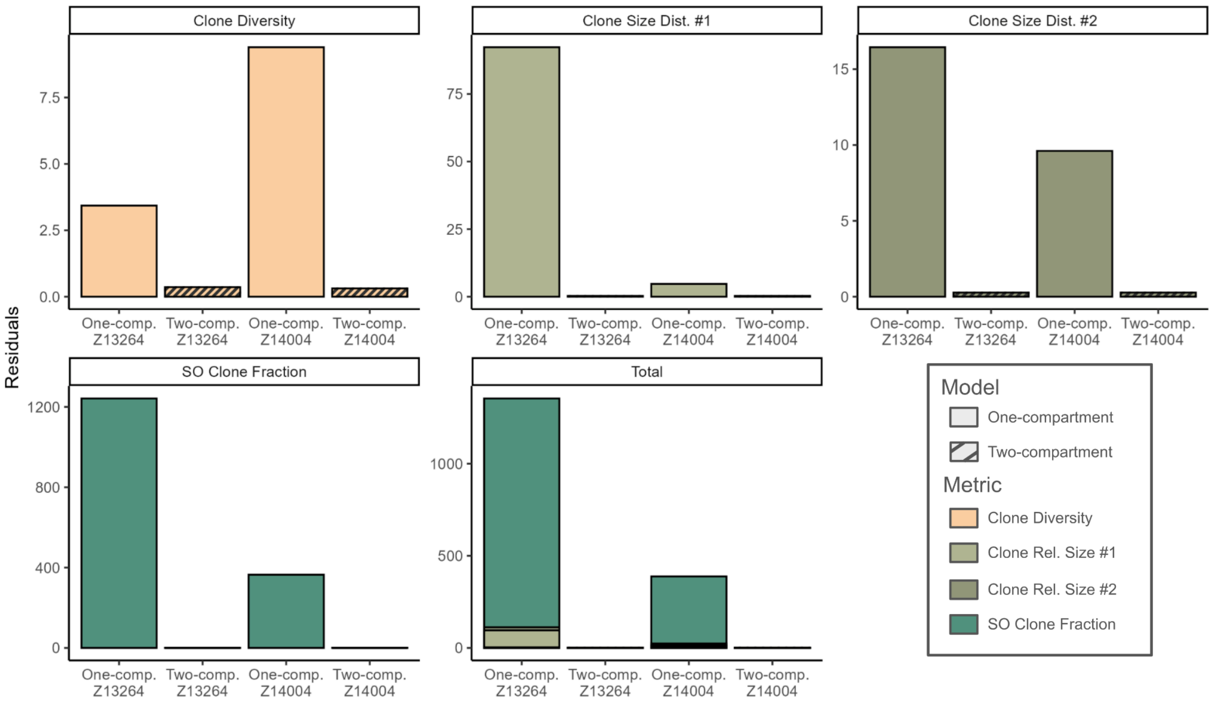


**Supplementary Figure S3:** Residual Sum of Squares (RSS) of best fits of one and two-compartment models optimized with respect to the metrics derived from data by Radtke et al (2023). Stacked bar plots show the RSS of each metric, diversity, SO clone fraction and clone size distribution in two different time points (see Supplementary Table 1), for two animals (Z13264 and Z14004). The two-compartment model provides consistently lower RSS across both animals and all clonal metrics, suggesting improved overall performance.

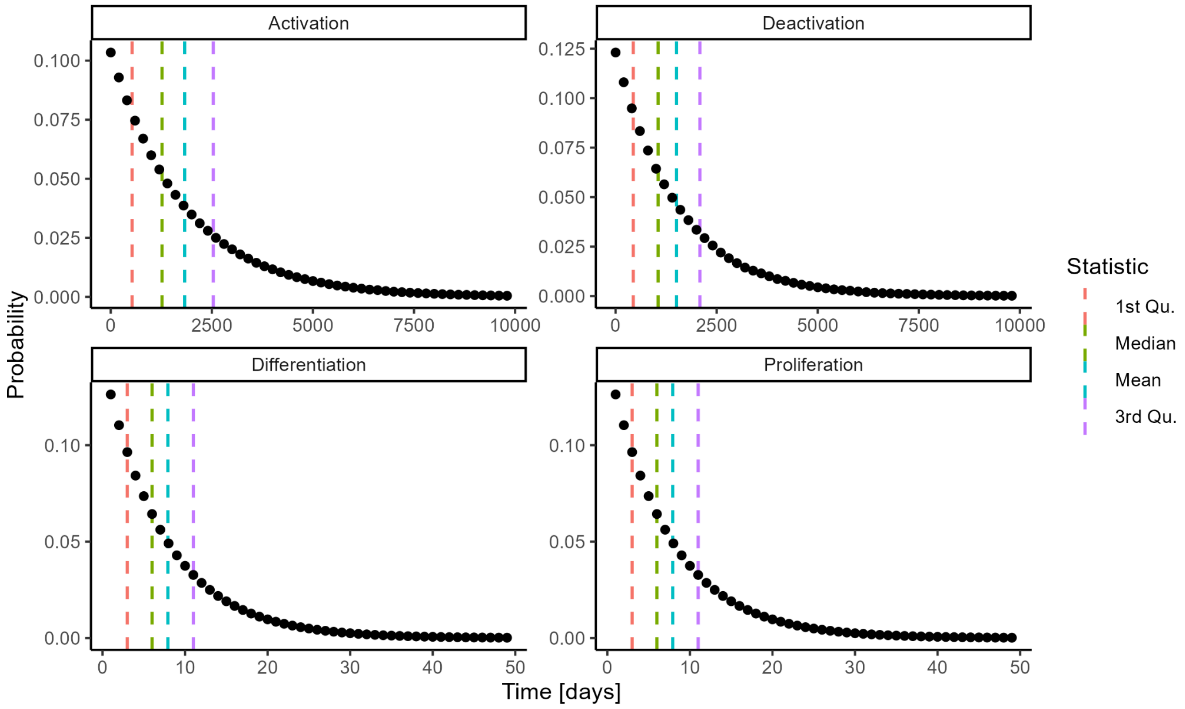

**Supplementary Figure S4:** Probability distributions of waiting times for individual cell events (activation, deactivation, differentiation, and proliferation) in the two-compartment model, estimated using best-fit parameters. Black dots indicate the probability that a single cell performs the indicated event after a given time. Vertical dashed lines show the 1st quartile (red), median (green), mean (blue), and 3rd quartile (purple) of each distribution, as summarized in Supplementary Table S4. Due to the exponential nature of these distributions, most events take place earlier than the mean waiting time, as shown by the 1st quartile and median statistics.

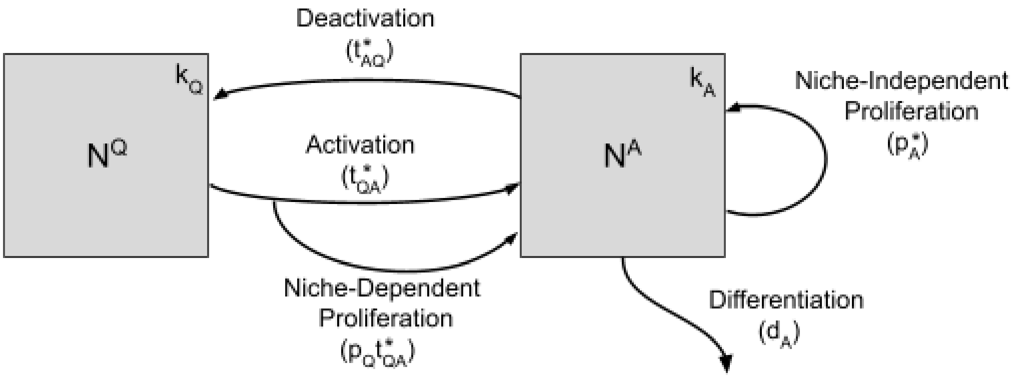


**Supplementary Figure S5:** Schematic representation of the unified model of HSCs dynamics. Gray boxes indicate the “active” and “quiescent” compartments, with carrying capacities $k_A$ and $k_Q$ and cell abundance $N^A$ and $N^Q$, respectively. Arrows indicate the flux of cells when the corresponding action is taken with propensities analogous to the ones defined on Table 1. The two-compartment model can be obtained by removing the niche-proliferation action.

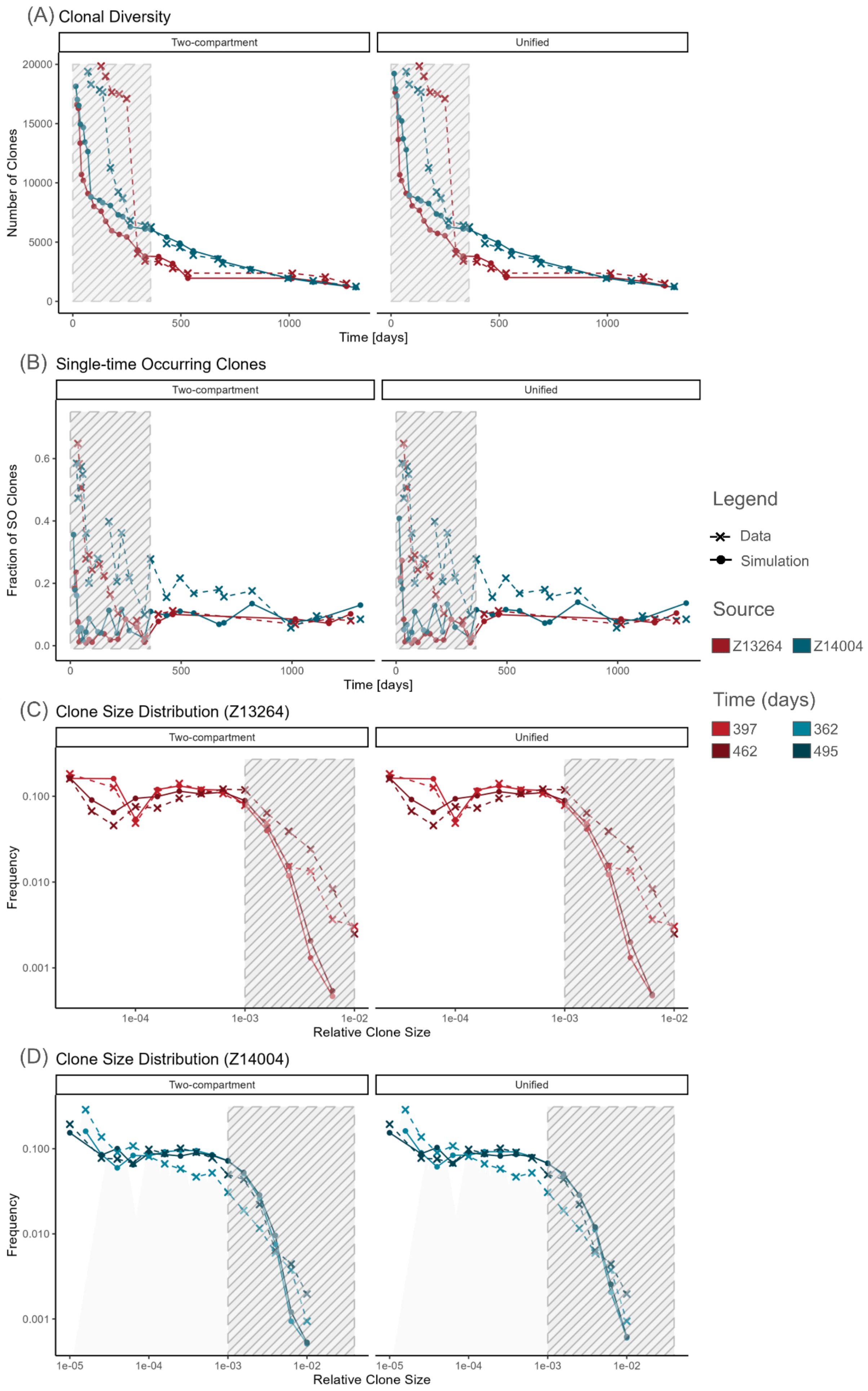

(A) Clonal Diversity
Two-compartment
Unified
Number of Clones
20000
15000
10000
5000
0
0
500
1000
Time [days]
(B) Single-time Occurring Clones
Fraction of SO Clones
0.6
0.4
0.2
0.0
Time [days]
(C) Clone Size Distribution (Z13264)
Frequency
0.100
0.010
0.001
1e-04
1e-03
1e-02
Relative Clone Size
(D) Clone Size Distribution (Z14004)
1e-05
Legend
Data
Simulation
Source
Z13264
Z14004
Time (days)
397
362
462
495

**Supplementary Figure S6:** Best fit for the two-compartment (A-D) and unified models (E-H) optimized with respect to the metrics derived from data by Radtke et al (2023). Dots indicate model predictions with lines connecting observations for clarity; experimental data is indicated by **x** marks. Striped areas indicate data which is not used for fitting.

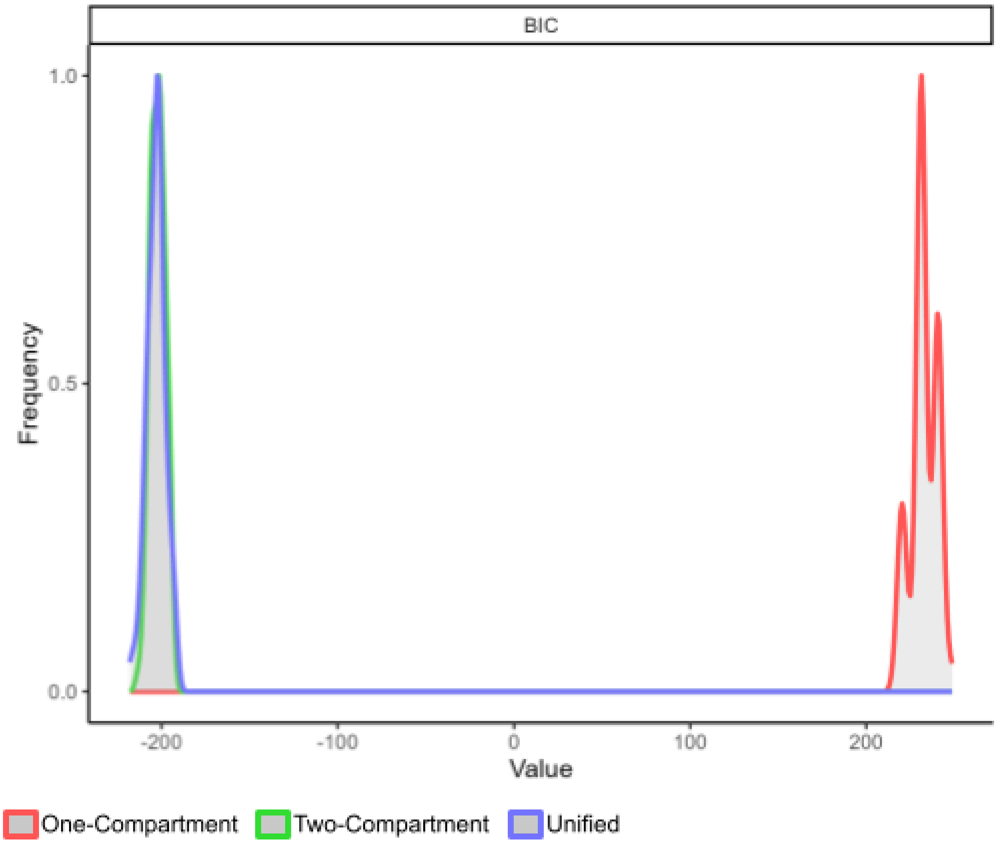


**Supplementary Figure S7:** Distribution of Bayesian Information Criteria (BIC) values from 100 simulations using the best-fit parameters of the one-compartment, two-compartment and unified models. Colors represent the different models, as in legend.

<table>
<tr><th></th><th>Z13264</th><th>Z14004</th></tr>
<tr><td rowspan="2">Diversity</td><td>Total Points: 21<br>Fitted Points: 6</td><td>Total Points: 25<br>Fitted Points: 10</td></tr>
<tr><td colspan="2">Criteria: Sampling day > 360.</td></tr>
<tr><td rowspan="2">SO Clone Contribution</td><td>Total Points: 21<br>Fitted Points: 5</td><td>Total Points: 25<br>Fitted Points: 10</td></tr>
<tr><td colspan="2">Criteria: Sampling day > 360 and exclude day 532 of Z13264.</td></tr>
<tr><td rowspan="3">Clone Size Distribution</td><td>Total Points: 235<br>Fitted Points: 17</td><td>Total Points: 313<br>Fitted Points: 20</td></tr>
<tr><td colspan="2">Criteria: Selected days and relative clone size <= $10^{-3}$.</td></tr>
<tr><td>Selected Days: 397 and 462</td><td>Selected Days: 362 and 495</td></tr>
<tr><td>Summary</td><td>Total Points: 277<br>Fitted Points: 28</td><td>Total Points: 373<br>Fitted Points: 40</td></tr>
</table>

**Supplementary Table S1:** Summary of data points used for model fitting across the three clonal aspects for animals Z13264 and Z14004. Diversity and SO clones fits were restricted to time points after homeostatic equilibrium, with SO clones further requiring over 100 reads. Clone size distribution was fitted with data on small and medium-sized clones from two early homeostatic-phase time points with high read coverage per animal.

| Parameter Name | Definition | Default Value |
|---|---|---|
| dA | Base differentiation rate for active cells. | - |
| pA | Base proliferation rate for active cells. | - |

| | | |
|---|---|---|
| tAQ | Base deactivation rate for active cells. | 0 |
| tQA | Base activation rate for quiescent cells. | 0 |
| clone_mult | Multiplicative factor for the number of clones.<br><br>Number of Clones = clone_mult*$10^4$ | - |
| pQ | Probability of a quiescent cell to proliferate when activating. | 0 |

**Supplementary Table S2:** Summary of parameters used in the optimization routine. The model rates (pA, dA, tAQ, tQA, and pQ) are directly optimized. The total number of clones is scaled by clone_mult. The model carrying capacities are kept fixed to $10^5$ cells on each compartment for two-compartment models and to 2*$10^5$ for one-compartment models. Default values are shown where applicable.

| Action | 1st Quartile | Median | Mean | 3rd Quartile |
|---|---|---|---|---|
| Activation | 527 | 1268 | 1828 | 2535 |
| Deactivation | 438 | 1051 | 1507 | 2085 |
| Differentiation | 3 | 6 | 8 | 11 |
| Proliferation | 3 | 6 | 8 | 11 |

**Supplementary Table S3:** Summary statistics of simulated waiting times for cell events in the two-compartment model. Estimates were derived from simulations run for 100,000 time steps using best-fit parameters of the two-compartment model.

| | **One Compartment** | **One Compartment (Downweighted SO)** | **Two Compartment** | **Unified** |
|---|---|---|---|---|
| Nr. Clones | 17.67*$10^4$ | 9.07*$10^4$ | 9.72*$10^4$ | 10.35*$10^4$ |
| $d_A$ | 0.045 | 0.088 | 0.126 | 0.108 |

| | | | | |
|---|---|---|---|---|
| $p_A$ | 0.435 | 0.620 | 0.708 | 0.494 |
| $k_A$ | $2*10^5$ | $2*10^5$ | $1*10^5$ | $1*10^5$ |
| $t_{AQ}$ | 0 | 0 | 0.061 | 0.055 |
| $t_{QA}$ | 0 | 0 | 0.003 | 0.002 |
| $k_Q$ | 0 | 0 | $1*10^5$ | $1*10^5$ |
| $p_Q$ | 0 | 0 | 0 | 0.054 |

**Supplementary Table S4:** Best fit parameters for four models of HSC clonal dynamics. Rates are determined by a customized R routine based on genetic algorithms while carrying capacities are kept fixed.
